# Production of Hyaluronan by the Trophectoderm is a Prerequisite for Mouse Blastocyst Attachment

**DOI:** 10.1101/2020.03.27.012880

**Authors:** Ron Hadas, Eran Gershon, Aviad Cohen, Michal Elbaz, Shifra Ben-Dor, Fortune Kohen, Nava Dekel, Michal Neeman

## Abstract

Embryo implantation requires execution of highly synchronized processes at the feto-maternal interface, initiated by blastocyst attachment to the endometrial epithelium. Hyaluronan is a major ECM component known to regulate adhesion-associated biological processes in various physiological settings. We hypothesized that hyaluronan may facilitate blastocyst attachment. In order to test our hypothesis, we characterized the blastocyst expression of hyaluronan synthesizing and degrading enzymes, as well as the expression of hyaluronan receptors during attachment. The functional impact of hyaluronan was challenged by the use of mouse transgenic blastocysts, in which genes encoding for hyaluronan synthesizing enzymes were deleted using lentiviral incorporation of Cas-9 endonuclease alongside specific short-guide RNAs into the embryonic trophectoderm. Embryos with transgenic trophectoderm were tested for their attachment *in vitro*, or assessed for implantation *in vivo*, upon transfer to foster dams. Deletion of the trophectoderm hyaluronan biosynthesis significantly reduced the number of blastocysts attached to human uterine epithelium cells *in vitro*. Reduced attachment was also observed *in vivo*, in pregnant mice carrying blastocysts with hyaluronan-depleted trophectoderm. In agreement, trophectoderm expression of osteopontin, was downregulated upon depletion of hyaluronan. MRI measurements revealed a decrease in uterine blood vessels permeability. Uterine expression of VEGF-A, PTGS-2 and uterine osteopontin, which constitute the immediate response to blastocyst attachment was also reduced. Furthermore, impaired implantation, associated with a decrease in hyaluronan synthesis in the mural trophectoderm, obtained upon tamoxifen treatment, has been recovered by LIF administration. These results demonstrate that estrogen-regulated hyaluronan-synthesis in the trophectoderm is indispensable for mouse blastocysts attachment to the uterine epithelium.

## Introduction

Implantation, during which the embryo attaches to and invades the maternal endometrium ^1^ is essential for successful pregnancy. In humans, natural conception per cycle is poor (~30%), and implantation defects were implicated in 75% of failed pregnancies ^2,3^. Implantation is initiated by blastocyst apposition, which in mice, takes place on the fourth day of pregnancy (E4.0), followed by the attachment of embryonic mural trophectoderm cells to the uterine luminal epithelium, and the subsequent invasion of the uterine epithelium into the implantation crypt toward the anti-mesometrial pole ^4^. These physiological events require both, a receptive uterus as well as a high quality blastocyst ^5^. Ovarian-secreted estrogen directly regulates blastocyst activation, in a temporally synchronized manner, preventing its dormancy ^6^. The activated blastocyst attaches to the uterine epithelium, stimulating in response, a local up-regulation of the pro-angiogenic factor, VEGF-A. A resulting increased endometrial vascular permeability in the implantation chamber has been monitored *in vivo* by dynamic contrast-enhanced (DCE) MRI, using a high-molecular weight contrast agent ^3,7,8^. Attachment and invasion are immediately followed by stromal cell decidualization, forming the primary decidual zone, consisting of PTGS-2 positive decidualized cells ^9^. Blastocyst attachment to the uterine anti-mesometrial epithelium is enabled by immediate cell surface interactions, mediated by extra-cellular matrix (ECM) carbohydrate contacts ^2^.

Hyaluronan is an anionic high-molecular weight polysaccharide, with an average molecular mass of 10^6^–10^7^ Da ^10,11^. Hyaluronan is produced in the inner side of the cell membrane by three hyaluronan synthases, HAS-1, HAS-2 and HAS-3. The production of hyaluronan by these membrane-bound enzymes is followed by the extrusion of its intact polysaccharide form, which is either retained in the cell surface or released ^12^. Hyaluronan can be subjected to extensive turnover by specific degrading enzymes known as hyaluronidases, the activity of which generates small oligosaccharides. The intact hyaluronan together with its shorter degradation products have been shown to directly regulate cell behavior via binding to specific hyaluronan receptors, such as CD44, RHAMM, LYVE-1 and TLR-4 ^13–15^.

The goal of this study was to determine the role of hyaluronan as a mediator of blastocyst attachment in mice. For that purpose, we generated blastocysts, in which the hyaluronan synthesizing enzymes were deleted using a Cas-9 endonuclease system (CRISPR). The use of lenti-viral vectors allowed exclusive targeting of these genetic modifications to the trophectoderm with no effect on the inner-cell mass (ICM) ^16^. Disruption of trophectoderm hyaluronan synthesis resulted in impaired attachment accompanied by poor decidual and angiogenic reaction detected in pregnant mice by MRI, as early as at E4.5. These results provide a novel addition to our knowledge of the ‘embryonic toolbox’ required for successful blastocyst attachment.

## Materials and methods

### Animals

C57Bl/6J female mice (6-12 week old; Envigo, Israel) were mated with Myr-Venus homozygote males. These hemizygote Myr-Venus embryos were used for histological analysis of hyaluronan metabolism at E4.5. Impairment of blastocyst activation *in vivo* was conducted as previously described ^5^. Briefly, pregnant ICR mice were intraperitoneally (i.p) administered at E2.5 with tamoxifen citrate (Sigma-Aldrich, Rehovot, Israel) 10 μg per mouse in a solution containing 12.5% ethanol, 12.5% cremophor EL (Sigma-Aldrich, Rehovot, Israel) and 75% of 5% dextrose, in water, which also served as vehicle. Recombinant mouse LIF (PeproTech, TX, USA) was administered i.p at a concentration of 10 ug in PBS/mouse at E3.5. Dams were sacrificed at E4.0 prior to dissection of uterine tissues. Uterine horns were subjected to embryo flushing using PBS, and immediately fixed in PFA 4% for immunohistochemical analysis. Flushed blastocysts were also fixed in PFA 4% prior to whole-mount immunofluorescence.

### Design and production of lentivectors

For the purpose of genomic interference with HAS-1 and HAS-2 expression, we have designed specific short guide RNA oligos (sgRNA) specific for the genomic site of interest (Table S1) The sgRNA sequences were cloned into lenti CRISPR v2 lentiviral vector, kindly provided by Dr. Igor Ulitsky (Weizmann Institute, Israel) according to a protocol previously described^17^ with slight modifications. Briefly, oligonucleotides for the HAS-1 or HAS-2 sgRNA guide sequences were phosphorylated using T4 PNK (NEB, Ipswich, MA) for 30 min at 37°C, then annealed by heating to 95°C for 5 minutes and cooled down to 25°C at 5°C/min. The lenti CRISPR v2 vector and the annealed oligos were then supplemented with FastDigest BsmBI (Thermo Fisher Fermentas, Waltham, MA) and T7 ligase (NEB, Ipswich, MA) by six cycles of 5 min at 37°C followed by 5 min at 23°C. The ligation reaction was next treated with PlasmidSafe exonuclease (NEB, Ipswich, MA) for 30 min at 37°C. Recombinant lentiviruses were produced by transient transfection in HEK293FT cells (Invitrogen, CA, USA), as described earlier ^18^, using 3 envelope and packaging plasmids and one of two viral constructs: (i) Lenti CRISPR v2 HAS-1 deletion (HAS-1 del), (ii) Lenti CRISPR v2 HAS-2 deletion (HAS-2 del), or (iii) Lenti CRISPR v2 (Control). Briefly, infectious lentiviruses were harvested at 48 and 72 hours post-transfection, filtered through 0.45-mm-pore cellulose acetate filters and concentrated by ultracentrifugation at 19400rpm, 15°C for 2.5 hours. Lentiviral supernatant titers were determined by Lenti-X p24 Rapid Titer Kit (supplemental Table 1) according to manufacturer’s protocol (Takara Bio USA, Inc. California, U.S.A).

### Mice and lentiviral transduction

ICR females (8-12 week-old) were mated with vasectomized males and assessed for the occurrence of vaginal plugs on the following day. Wild-type ICR females (3-4 weeks old) were super-ovulated by sub-cutaneous injection of pregnant mare’s serum gonadotropin (PMSG) (Sigma-Aldrich, Rehovot, Israel) (5 units) followed 48 hours later by i.p injection of human chorionic gonadotropin (hCG) (Sigma-Aldrich, Rehovot, Israel) (5 units) and then mated with wild-type ICR males. Morulae stage embryos were collected from the females at E2.5 and then incubated in KSOM medium to obtain expanded blastocysts. Following removal of the zona pellucida by Tyrode’s solution (Sigma-Aldrich, Rehovot, Israel) embryos were incubated with lentiviruses, in KSOM for 5 hours. The transduced blastocysts implanted into pseudo-pregnant ICR uteri generated after mating with vasectomized ICR males by non-surgical embryo transfer (NSET, ParaTechs, Kentucky, US). All mice were maintained under specific pathogen–free conditions and handled under protocols approved by the Weizmann Institute Animal Care and Use Committee according to international guidelines.

### In vitro blastocyst attachment assay

Blastocyst attachment assay was performed as described previously ^19,20^. Briefly, blastocysts were infected with lentiviruses for five hours, washed and immediately labeled with Vybrant Cell-Labeling Solution (ThermoFisher Scientific, Waltham, MS) for 20 minutes, before transferring unselectively to confluent Ishikawa cell monolayers in a 96-well plate coated with Matrigel (Invitro technologies, Victoria, Australia). Following florescent labeling, a total of three blastocysts was transferred per well. Co-cultures were incubated undisturbed at 37 °C in a 5% CO2 atmosphere for 48 hours. The stability of embryo attachment was measured by repeated aggressive washing and tilting of the culture plates. Attached embryos were detected using a fluorescent stereo microscope (Nikon, Tokyo, Japan). The ratio of attached embryos per well was defined as a single observation.

### In vivo Dynamic Contrast–Enhanced (DCE) MRI of embryo implantation sites

DCE MRI experiments were performed at 9.4 T on a horizontal-bore Biospec spectrometer (Bruker, Karlsruhe, Germany). The pregnant mice were serially scanned after transgenic embryo transplantation at E4.5. Three-dimensional gradient echo (3D-GE) imaging of embryo implantation sites was conducted as previously described ^7^ yielding two parameters that characterize vascular development and function: (i) fractional Blood Volume (fBV), which describes blood-vessel density, and (ii) Permeability Surface area product (PS), which represents the rate of contrast agent extravasation from blood vessels and its accumulation in the interstitial space, Mean fBV and PS were calculated separately for single implantation sites considering homogeneity of variances between mice.

### Immunohistochemistry analysis

Uterine sections containing embryo implantation sites were fixed in 4% paraformaldehyde (PFA) and embedded in paraffin. For morphological analysis, tissues were either stained with hematoxylin and eosin, or underwent immunohistochemical stainings. Blastocysts were fixed in 4% PFA and underwent immunofluorescence according to standard protocols, with the following primary antibodies: goat anti HAS-1 (sc-23145; Santa Cruz Biotechnology, Dallas, TX, USA) goat anti HAS-2 (sc-34067; Santa Cruz Biotechnology), mouse anti TLR-4 (sc-293072; Santa Cruz Biotechnology), goat anti GFP (ab6673; Abcam), rabbit anti CD44 (ab41478; Abcam), rabbit anti RHAMM (ab124729; abcam), rabbit anti LYVE-1 (70R-LR003; Fitzgerald Industries International, Acton, MA, USA), Rabbit anti osteopontin (NBP1-59190; Novus biologicals, CO, USA), Rabbit anti VEGF (sc-152; Santa Cruz Biotechnology) and Sheep anti hyaluronan (ab53842, Abcam), Rabbit anti PTGS-2 (Cayman chemical, Michigan, USA), Rabbit anti Hyal-2 (ab68608, Abcam), Rabbit anti Hyal-1(ab203293, Abcam). Next, slides were incubated with secondary antibodies, conjugated to biotin against the appropriate species (except for goat anti GFP in the case of double staining. and incubated with fluorophore conjugated StreptAvidin (Jackson Immunoresearch Laboratories, PA, USA), Cells undergoing apoptosis were detected by TUNEL staining (ApopTag; Merck Millipore, MA USA). Slides were counterstained with hematoxylin and subsequently mounted. All slides were imaged using a fluorescent Olympus SZX-RFL2 zoom stereo microscope. Flushed blastocysts were fixed with PFA 4% for 20 minutes, washed in PBST 0.1% and permeabilized by 0.5% Triton-X100 for 15 minutes, prior to their incubation with primary antibodies in PBS containing 3% bovine serum albumin, overnight at 4C. After washes, the blastocysts were incubated for two hours with secondary antibodies against appropriate hosts, conjugated to alexa-fluor fluorophores (1:250 dilution in PBS), counterstained with Hoechst (Invitrogen) and subsequently mounted in paraffin oil. Images were then acquired using a Zeiss LSM710 confocal microscope and spinning disk 386 confocal microscope (Zeiss, Oberkochen, Germany) and quantified by FIJI software (https://imagej.nih.gov/ij/).

### Statistical Analysis

All statistical analyses performed in this study were two-tailed with a similar level of significance (p=0.05), Tukey Kramer’s post-hoc significance test and demonstrated normal values distribution. On-way ANOVA analysis of variance was conducted for: immunofluorescent detection of hyaluronan production following tamoxifen and mLIF treatment before implantation (**Figure 3**), Blastocyst attachment assays (**Figure 5**) and DCE-MRI of pregnant mice (**Supplementary figure 2 and Figure 7**). All statistical analyses were conducted using GraphPad Prism 7(CA, USA).

## Results

During the final stages of pre-implantation development, the blastocyst undergoes lineage differentiation, generating the polar trophectoderm, adjacent to the ICM, and the mural trophectoderm that attaches to the uterine epithelium and initiates the invasion to the underlying stroma. Both HAS-1 and HAS-2 are prominently expressed at E4.5, in the trophectoderm during blastocyst growth and expansion (**Fig. 1a-b**) as well as in the attachment sites (**Fig. 1c-d**), whereas the expression of Has-3 was hardly detected. This correlates with accumulation of hyaluronan in the trophectoderm observed during attachment (**Fig. 1e-f & Fig. S1a-b**). The hyaluronan degrading enzymes. Hyal-1 and Hyal-2, were detected in the luminal epithelium at the attachment sites, with hyal-2 also observed in the trophectoderm **(Fig. S1c)**. Expression of hyaluronan receptor, CD44, was observed in blastocysts as well as in uterine luminal epithelium, prior to attachment (**Fig. 2a-b**). Nevertheless, another ligand of CD44, osteopontin was also identified in the trophectoderm of late blastocysts **(Fig. S1f)**. Other hyaluronan receptors, such as RHAMM and LYVE-1 (**Fig. 2c-d**) were detected at E4.5 in the luminal epithelium, specifically in attachment sites, as well as in the sub-epithelial stroma, while TLR-4 was upregulated in the trophectoderm (**Fig. 2e-f**). Altogether, we observed the hyaluronan production enzymes in the trophectoderm, alongside its degrading enzymes as well as the corresponding receptors in the adjacent uterine epithelium.

**Figure 1.**
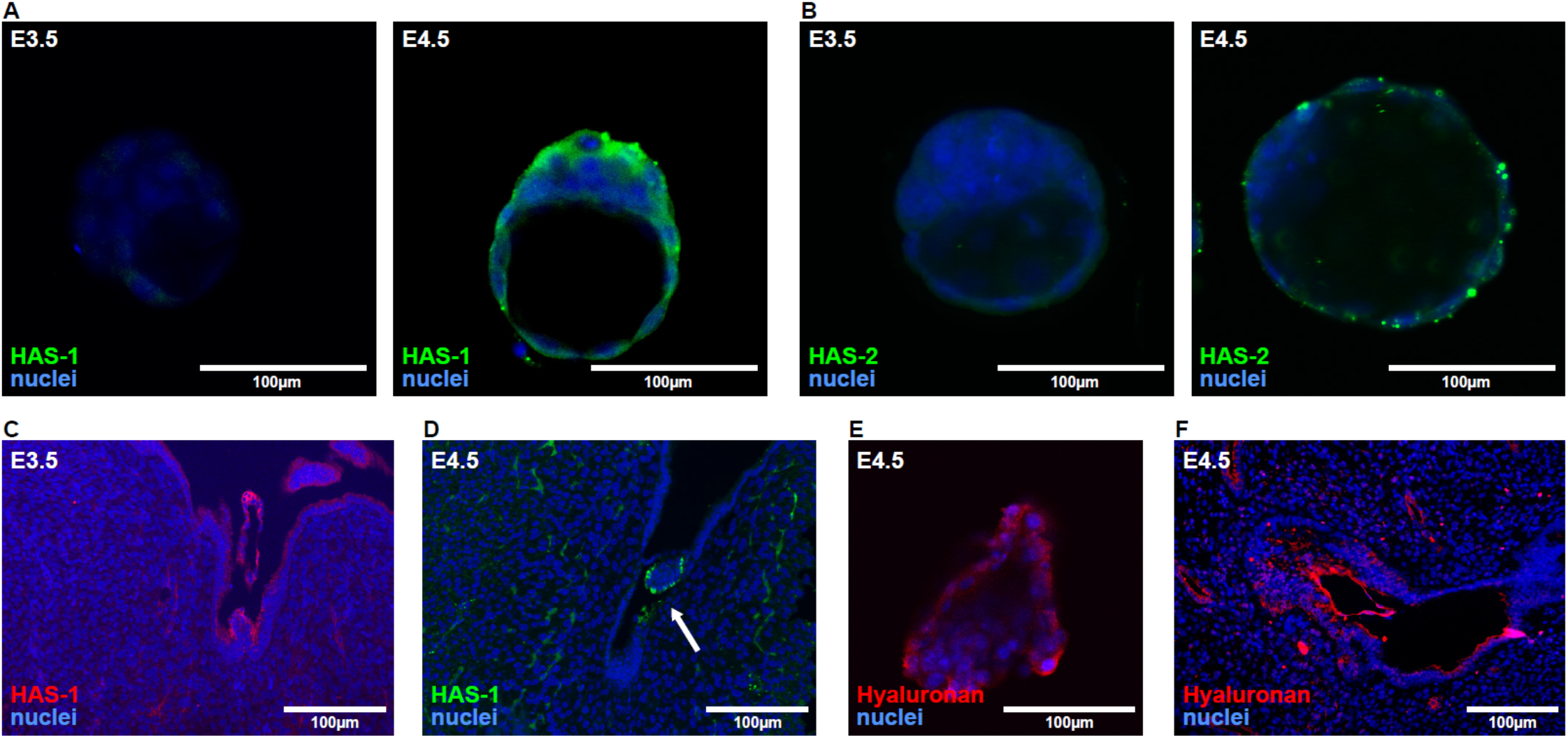
Hyaluronan synthesis and accumulation in the attachment interface. Female mice were mated with Venus+ males, their uterine horns were harvested at E3.5 and E4.5 and immediately flushed. (n=3 dams). (**A**) Immunofluorescent staining of HAS-1 in blastocysts. (**B**) Immunofluorescent staining of HAS-2 in blastocysts. (**C**) Immunohistochemical detection of HAS-1 in histological sections. (**D**) Immunohistochemical detection of HAS-2 in histological sections; embryo indicated by a white arrow. (**E**) Immunofluorescent staining of hyaluronan in blastocysts. (**F**) Immunofluorescent staining of hyaluronan in peri-implantation blastocysts. (**E**) Immunohistochemical detection of hyaluronan in histological sections.

**Figure 2.**
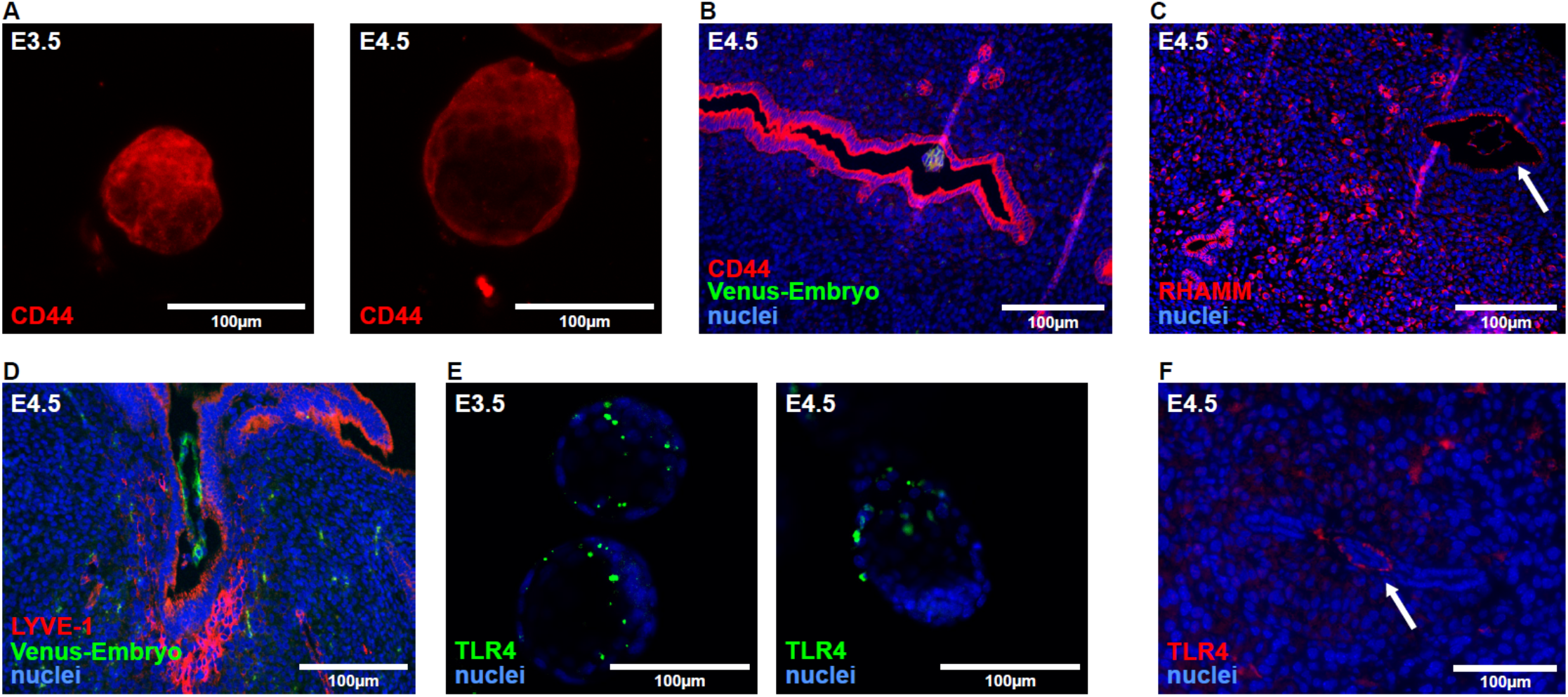
Hyaluronan receptors in the attachment interface. Female mice were mated with Venus+ males, their uterine horns were harvested at E3.5 and E4.5 and immediately flushed. (n=3 dams). (**A**) Immunofluorescence staining of CD44 in blastocysts. (**B**) Immunohistochemical staining of CD44 in histological sections. (**C**) Immunohistochemical detection of RHAMM in histological sections; embryo is indicated by a white arrow. (**D**) Immunohistochemical detection of LYVE-1 in histological sections. (**E**) Immunofluorescent staining of TLR-4 in blastocysts. (**F**) Immunohistochemical detection of TLR-4 in histological sections; embryo is indicated by a white arrow.

To test whether hyaluronan production by the trophectoderm during blastocyst activation is regulated by estrogen, the estrogen competitor, tamoxifen, was injected to pregnant mice at E2.5. A decrease in estrogen-regulated osteopontin expression in the glandular epithelium at E4.0 validated the effect of tamoxifen (**Fig. 3a**). Blastocysts flushed at E4.0 from tamoxifen treated mice showed a reduced HAS-1 expression in the trophectoderm, resulting in a corresponding decrease in hyaluronan accumulation on the surface of the trophectoderm. This effect was reversed by administration of the estrogen downstream effector, LIF at E3.5 (**Fig. 3b-e**). In sum, we revealed hyaluronan deposition by the trophectoderm to be downstream of the ovarian estrogen nidatory surge, mediated by uterine secretion of LIF.

**Figure 3.**
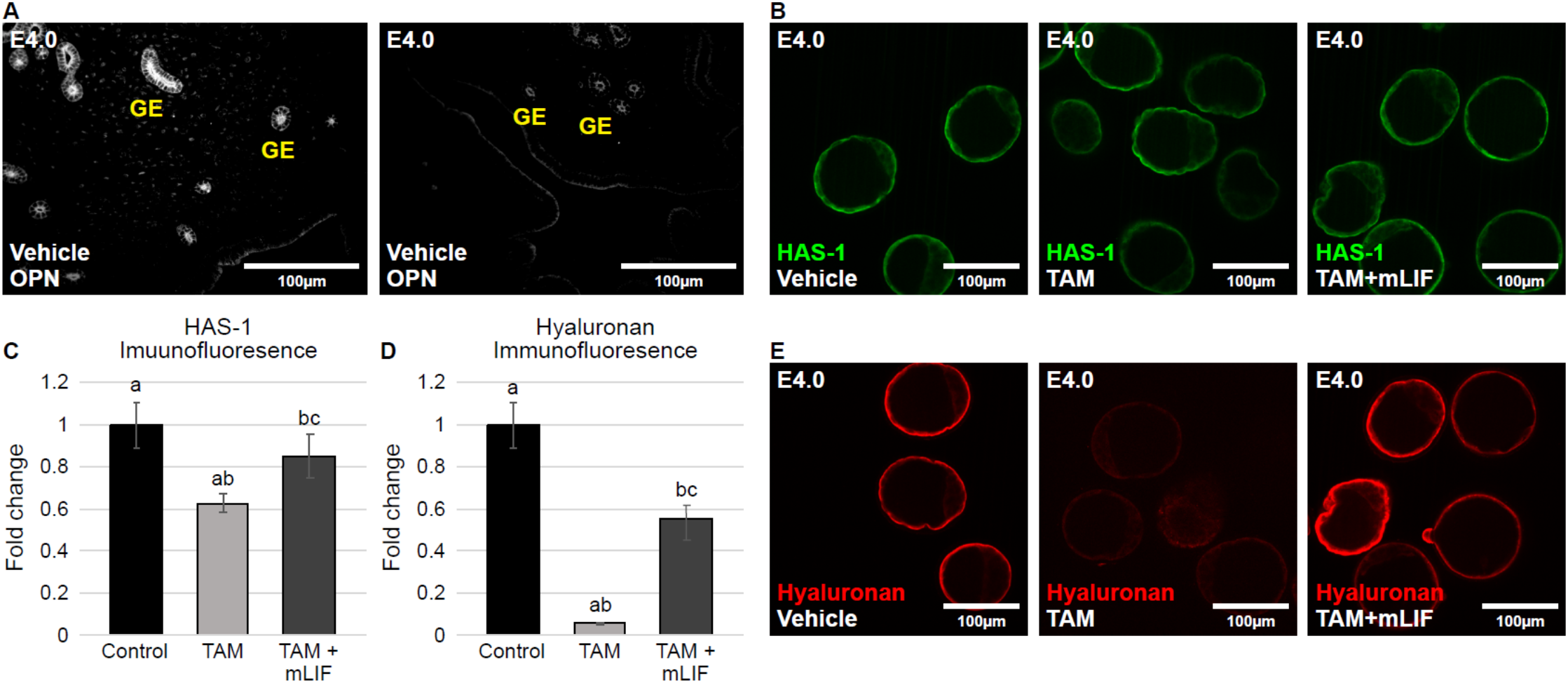
Glandular secretion of LIF-regulated estrogen, triggers hyaluronan synthesis in the trophectoderm. ICR female mice mated with ICR males, were administered with tamoxifen at E2.5, followed by recombinant mouse LIF; their uterine horns were harvested at E4.0 and immediately flushed (n=3 dams). (**A**) Model validation by immunohistochemistry of osteopontin (OPN) in E4.0 uteri following tamoxifen (TAM) treatment; GE indicates glandular epithelium. (**B**) Immunofluorescent staining of HAS-1 in blastocysts following tamoxifen treatment and recombinant mouse LIF (mLIF). (**C–D**) Image analysis of HAS-1 and hyaluronan in the trophectoderm following tamoxifen treatment and mouse LIF repletion; Letters indicate statistically significant differences (±0.108; 0.041; 0.105) (n=3 dams, 8 embryos in Vehicle group, 19 embryos in TAM, 9 embryos in mLIF); (±0.109; 0.005; 0.07) (n=3 dams, 8 embryos in Vehicle group, 19 embryos in TAM, 9 embryos in mLIF);. (**E**) Immunofluorescent staining of hyaluronan in blastocysts following tamoxifen treatment and mLIF injection.

The role of maternal hyaluronan during implantation was examined by pharmacological suppression of hyaluronan synthesis, using DON (1μg/g BW; intraperitoneally administered daily to pregnant mice), an effective inhibitor of glucosamine synthesis ^21^. DON-treated mice showed similar number of implantation sites as controls, PBS-treated mice (**Fig. S2a)**, ruling out a role for maternal hyaluronan on this early stage of pregnancy.

Abrogation of hyaluronan synthesis targeted to the trophectoderm was achieved by a CRISPR Cas-9 endonuclease system, using lenti-viral delivery. As expression of Has3 was hardly detected at E4.5 deciduae specific guides exclusively against Has1 and Has2 were employed (**Fig 4a**). Immunofluorescence of blastocysts, showing decreased expression strictly in the trophectoderm (**Fig 4b-c**), as well as in implantation sites harvested from foster dams at E4.5 (**Fig 4d-e**) validated this model. *In vitro* attachment was examined by superimposing these transgenic blastocysts, previously incubated with a vital dye (**Fig. 5a**), on the surface of uterine epithelial cell line^20^. Trophectoderm deletion of either Has1 or Has2, and most remarkably of both, reduced blastocyst attachment *in vitro* (**Fig. 5b**). To examine the role of trophectoderm-synthesized hyaluronan *in vivo*, the transgenic embryos were transferred to pseudo-pregnant mice, serving as surrogate mothers. The uteri of these mothers were then flushed at E4.75, a time at which, in ICR (CD1) mice, stable blastocyst attachment should have been accomplished. Double-deletion of Has1 and Has2 resulted in reduced blastocyst attachment, indicated by an increased number of embryos successfully flushed out of the uterine lumen (**Fig. 5c**). All in all, targeted deletion of hyaluronan synthesizing enzymes in the embryo’s trophectoderm prevented effective blastocyst attachment.

**Figure 4.**
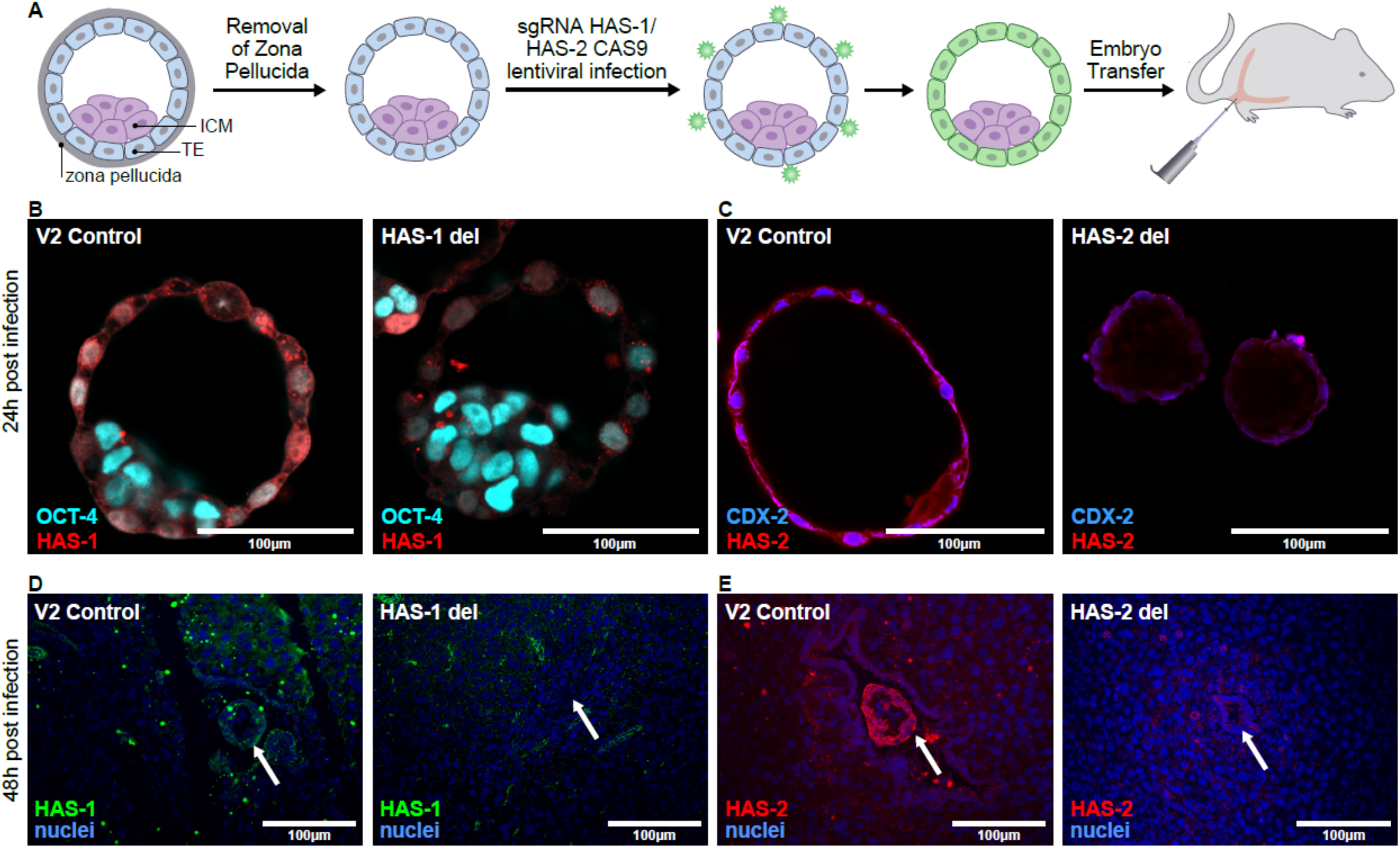
Hyaluronan depletion in the trophectoderm by CRISPR implementation using lenti-viral infection. Pre-implantation embryos flushed from ICR female mice subjected to lenti-viral infection of the trophectoderm was followed by embryo transfer to pseudo-pregnant mice. (**A**) Illustration of CRISPR-mediated interference with peri-implantation trophectoderm synthesis of hyaluronan. (**B**) Immunofluorescent staining of HAS-1 and Oct-4^+^ ICM cells in blastocysts grown in culture for 24 hours after viral infection. (**C**) Immunofluorescent staining of HAS-2 and CDX-2^+^ trophectoderm in blastocysts grown in culture for 24 hours after viral infection. (**D-E**) Immunohistochemical detection of HAS-1 and HAS-2 in E4.5 deciduae of foster dams.

**Figure 5.**
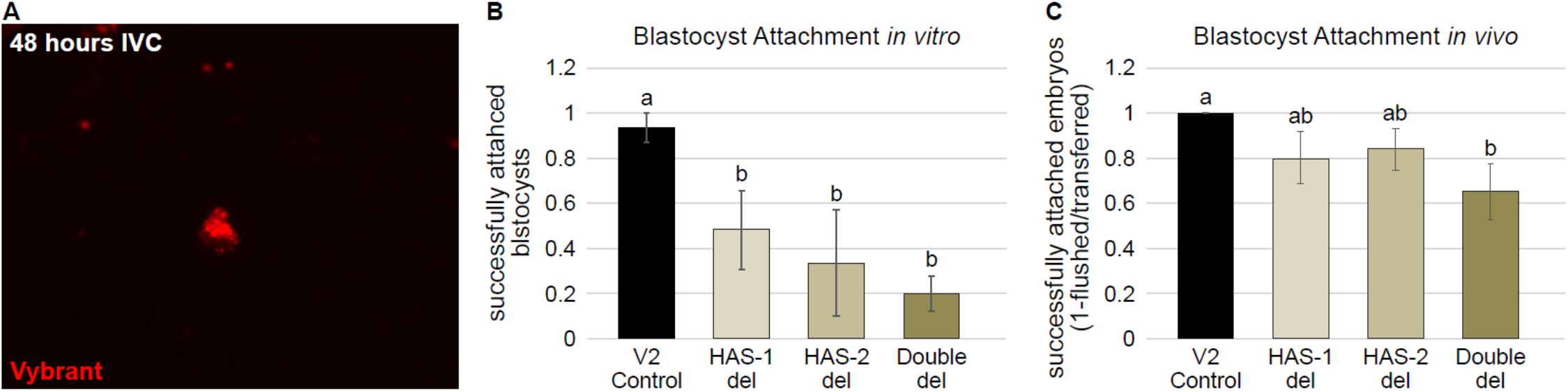
Trophectoderm production of hyaluronan is essential for blastocyst attachment. Pre-implantation embryos, subjected to lenti-viral infection of the trophectoderm, were flushed from ICR Female mice. These embryos were either cultured *in vitro* (IVC) with an endometrial carcinoma cell line, or transferred to pseudo-pregnant mice. (**A**) Longitudinal detection of blastocysts during IVC was achieved by their labeling with Vybrant Cell-Labeling Solution. (**B**) Quantification of the *in vitro* attachment assay, at 48 hours; letters indicate statistically significant differences (±0.066; 0.175; 0.235; 0.081) (n=15 embryos per group). (**C**) Quantification of *in vivo* attachment assay conducted in foster dams at E4.75; letters indicate statistically significant differences (±0; 0.1154; 0.092; 0.125) (n=5 dams in Control, 4 dams in HAS-1 del, 5 dams in HAS-2 del, 4 dams in Double del).

Successful implantation of blastocysts in the control group was visualized by H&E staining of embryo deciduae harvested from surrogate mothers (**Fig. 6a**). TUNEL assay detected prominent embryonic cell death in the treated groups, most prominently in embryos with dual deletion of hyaluronan synthesizing enzymes (**Fig. 6b**). The latter was associated with a decrease in the expression of osteopontin, previously demonstrated to correspond with primary uterine decidualization^22^, at the attachment interface (**Fig. 6c**). To test the key vascular response that accompanies uterine decidualization in mice, we assessed local uterine vascular function, following blastocyst attachment of transgenic embryos. For that purpose, we performed DCE MRI of foster dams at E4.5. We found that the deletion of hyaluronan synthesizing enzymes attenuated angiogenesis and reduced fractional blood volume (**Fig. 7a-d**). Impaired vascular response was confirmed by examination of histological sections, in which the distribution of the injected contrast agent biotin-BSA-Gd-DTPA was compromised (**Fig 7e**), alongside a reduced expression in the attachment interface, of VEGF-A, which serves as a potent decidual pro-angiogenic factor in response to blastocyst apposition (**Fig 7f**). Aberrant decidual response, as a result of compromised hyaluronan synthesis, was also demonstrated by decreased PTGS-2 in primary decidualized cells (**Fig. 7g**).

**Figure 6.**
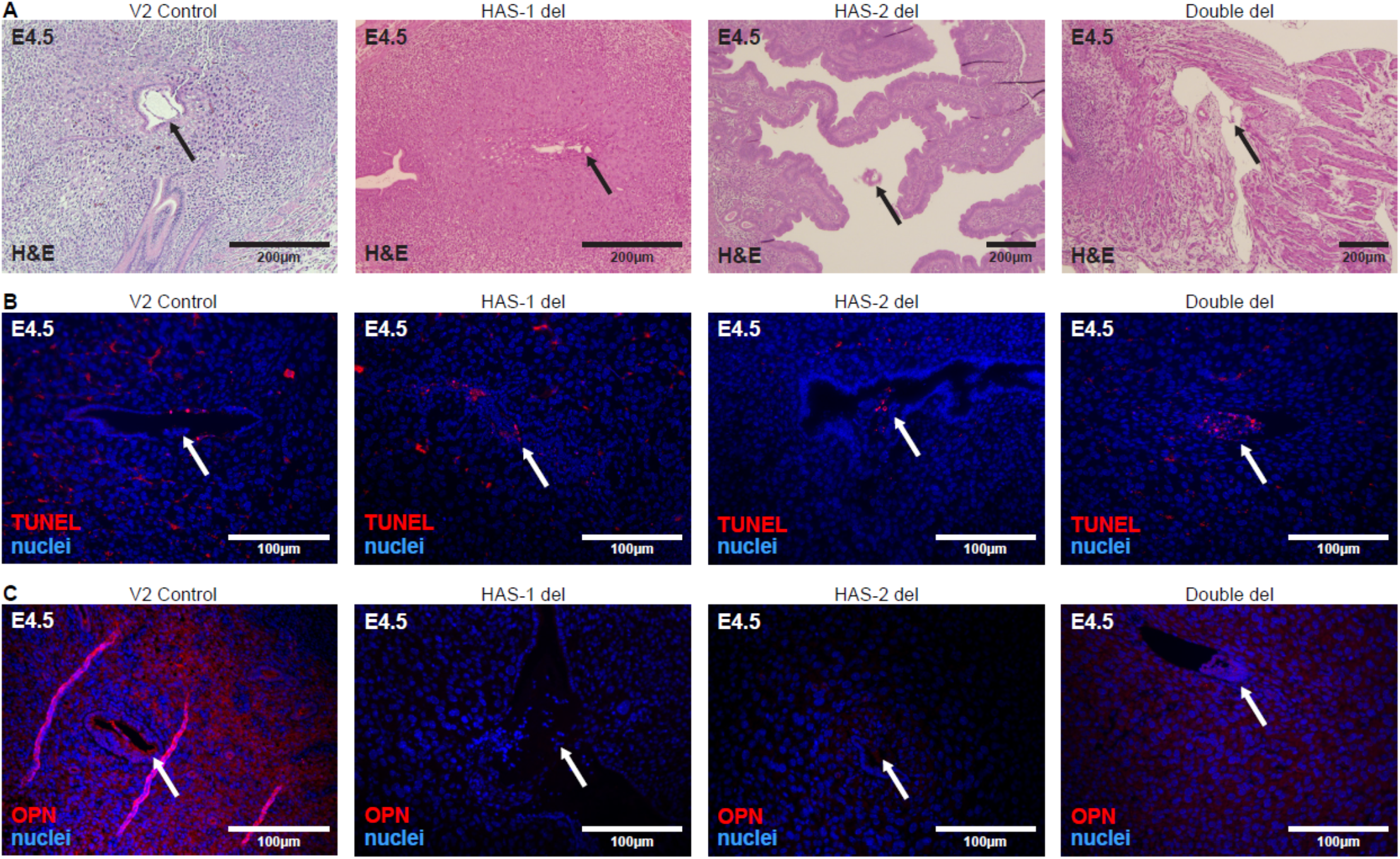
Hyaluronan depleted blastocysts fail to implant. Deciduae were harvested from foster dams of all groups at E4.5, immediately fixed and subjected to histological analysis. (**A**) H&E staining of embryo implantation sites; black arrows indicate blastocysts. (**B**) TUNEL analysis of E4.5 deciduae; white arrows indicate embryos. (**C**) Immunohistochemical detection of osteopontin (OPN) in E4.5 deciduae.

**Figure 7.**
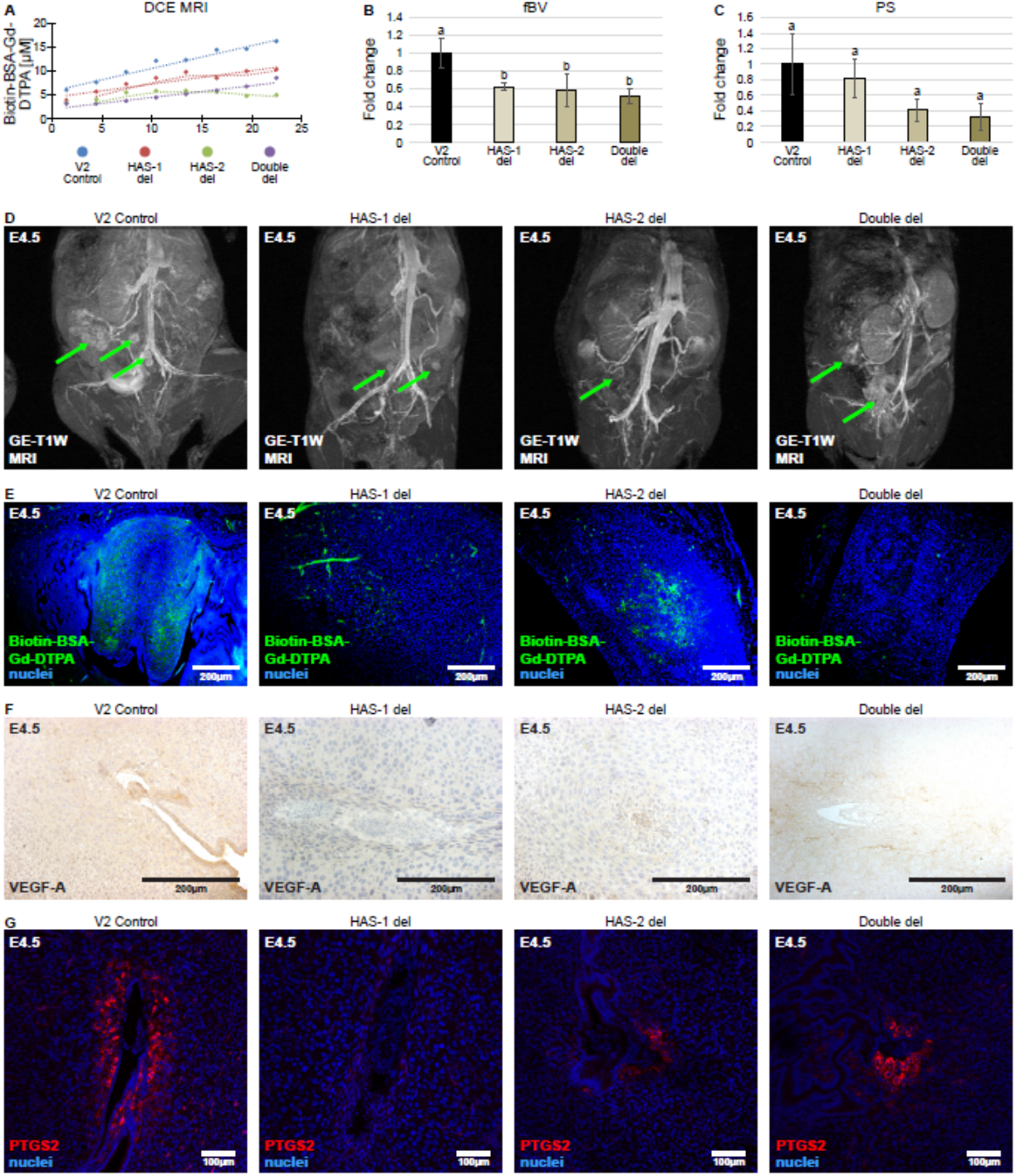
Trophectoderm synthesized hyaluronan regulates the primary decidual response. Foster dams of all groups were subjected to DCE MRI at E4.5; their deciduae were harvested, fixed and subjected to histological analysis. (**A**) Linear regression plots generated by DCE MRI analysis of all four groups; letters indicate statistically significant differences. Two functional vascular parameters produced from DCE MRI analysis. (**B)** Fractional blood volume (fBV); (±0.199; 0.039; 0.184; 0.093) (n=8 deciduae in Control,8 deciduae in HAS-1 del, 5 deciduae in HAS-2 del, 5 deciduae in Double del), (**C**) and permeability surface area (PS) respectively; letters indicate statistically significant differences (±0.384; 0.24; 0.14; 0.171) (n=8 deciduae in control, 8 deciduae in HAS-1 del, 5 deciduae in HAS-2 del, 5 deciduae in Double del). (**D**) Representative maximum intensity projection images of foster dams acquired during T1-weighted MRI. (**E**) *Ex-vivo* detection of intravenously administered MRI contrast agent, 30 minutes after injection of E4.5 deciduae; white arrows indicate embryos. (**F**) Immunohistochemical analysis of VEGF-A in implantation sites. (**G**) Immunohistochemical analysis of PTGS-2 in implantation sites.

## Discussion

The goal of this study was to investigate the significance of hyaluronan in blastocyst attachment and successful gestation. Immunofluorescent analysis of pre-implantation embryos, inhibition of estrogen receptor signaling and lineage specific genomic interference with hyaluronan synthesis, all demonstrated the essential role of hyaluronan metabolism within the chain of events preceding embryo implantation. Specifically, our study deciphered the crucial role of estrogen-regulated trophectoderm-synthesized hyaluronan for blastocyst attachment, decidualization and the primary maternal vascular response.

Considering the classical role of hyaluronan as a biological glue, its possible role in facilitating attachment of the pre-implantation embryo has been hypothesized. The expression dynamics of hyaluronan and its receptors and its accumulation in the uterine-blastocyst interface demonstrated previously by others ^5,6,23^, and shown herein by us, reinforced the notion that this ECM component may indeed mediate blastocyst adhesion. This idea was further supported by hyaluronan synthesis and subsequent accumulation prior to implantation. Interestingly, multiple key hyaluronan receptors were detected in the implantation chamber; TLR-4 and CD44 were observed in the attachment interface, and LYVE-1 and RHAMM were detected in the underlying stroma. Alongside with their biochemical capacity to bind hyaluronan, these receptors could potentially bind hyaluronan oligosaccharides generated by locally distributed hyaluronidases, to facilitate trophoblast invasion, when the intact hyaluronan molecule might interfere ^24,25^. The most prominent receptor for hyaluronan, CD44, was detected in the uterine epithelium at E4.5. Interestingly, CD44 is also a receptor for osteopontin, a secretory product of uterine glandular epithelium, expressed in response to the acute increase in ovarian estrogen, responsible for blastocyst activation in mice ^5,6^. Osteopontin was also observed in the trophectoderm of flushed embryos. Therefore, CD44 could also serve for osteopontin binding. Previous studies reported that depletion of CD44 in the rabbit resulted in impaired embryo implantation, whereas ablated CD44 in mice did not affect litter size ^26^. Furthermore, hyaluronan-CD44 interaction, was demonstrated to be employed by the embryo and uterine cells *in vitro* ^27^. These conflicting phenotypes may imply the redundant character of hyaluronan metabolism and receptor binding as previously demonstrated ^12,28^.

Secretion of LIF and its downstream target osteopontin by the glandular epithelium is regulated by synchronized estrogen-estrogen receptor signaling and sequentially obligatory for epithelial receptivity and primary decidualization via COX-2 activity ^5,6,29^. Our study demonstrates that hyaluronan synthesis is also regulated by estrogen. This conclusion is supported by the effect of the estrogen competitor, tamoxifen ^5^, in reducing deposition of hyaluronan in the trophectoderm associated with decreased HAS-1 expression. Phenotype rescue by pre-implantation supplementation of LIF further positions hyaluronan synthesis as a downstream target of glandular epithelial, estrogen-stimulated, LIF secretion, within the chain of events leading to blastocyst activation. Nevertheless, we cannot rule out a direct regulation of hyaluronan deposition by embryonic estrogen receptor, expressed by murine mural trophectoderm ^30^.

The fact that hyaluronan is deposited at the maternal-embryo interface and that this event is subjected to estrogen regulation may point indeed to its role in implantation. Nevertheless, the origin of the deposited hyaluronan remained to be identified. It should be noted that at the attachment interface, HAS-1 and HAS-2 are expressed mainly by the trophectoderm. Nevertheless, we examined the impact of metabolic suppression of maternal hyaluronan synthesis on embryo implantation. The production of maternal hyaluronan was inhibited by DON, which prevents the biosynthesis of glucosamine-6-P, an intermediate metabolite in the synthesis of UDP-N-acetyl glucosamine, a substrate of the three HAS isozymes ^31^. The fact that pre-treatment with DON did not alter the number of implanted embryos implies that uterine hyaluronan is not required for blastocyst attachment. To provide further support for the embryonic origin of hyaluronan we used mouse transgenic blastocysts, in which genes encoding for hyaluronan synthesizing enzymes were deleted using lentiviral incorporation to the embryonic trophectoderm. Embryos, in which hyaluronan synthesis in the trophectoderm was suppressed failed to attach. The most robust phenotype was observed in embryos in which both hyaluronan synthases were deleted. The latter also demonstrated decreased osteopontin expression by decidual cells as well as by attached embryos. These findings are consistent with the increased osteopontin levels observed in endometrial cells next to attached blastocysts ^20^, the identification of osteopontin as a transcriptional target of hyaluronan ^32^ and trophectoderm production of osteopontin as a target of ovarian secreted estrogen ^33^. The requirement for double-deletion of both hyaluronan synthases to prominently obtain embryonic mortality, impaired blastocyst attachment and poor decidual morphology, suggests redundancy of HAS-1 and HAS-2 in hyaluronan metabolism. Defective primary decidualization was also reflected by decreased PTGS-2 expression by peri-embryo decidualized cells, which is typically initiated at E4.25 adjacent to the implanting blastocysts, forming the primary decidual zone ^9^. Furthermore, the attenuated vascular response in the absence of embryonic hyaluronan, revealed by functional

MRI inspections, accompanied by a reduced trophectoderm expression of VEGF-A, might result either from weak blastocyst apposition or decreased PTGS-2 in decidualized stroma, as previously demonstrated post-implantation in PTGS-2 null mice ^34^.

In summary, our study demonstrates that hyaluronan production by the trophectoderm is a synchronized, hormonally regulated process, indispensable for successful blastocyst attachment and primary decidualization in mice. The present protocols of *in vitro* fertilization treatments call for single-embryo transfer, thus raising the need for identification of molecular markers for the selection of the high-quality embryo. Our study suggests that the capacity for hyaluronan biosynthesis may possibly serve for ranking the attachment capacity of blastocysts.

## Acknowledgement

This work was supported by the Seventh Framework European Research Council Advanced Grant 232640-IMAGO and by National Institutes of Health (grant 1R01HD086323-01). Michal Neeman is incumbent of the Helen and Morris Mauerberger Chair in Biological Sciences.

## Author contributions

R.H designed research studies, conducted experiments, acquired data, analyzed data, and wrote the manuscript. E.G designed research studies and conducted experiments. A.C designed research studies and conducted experiments. M.E conducted experiments. S.B.D designed short-guide RNAs. F.K designed research studies and reviewed the manuscript. N.D designed research studies and wrote the manuscript. M.N designed research studies and wrote the manuscript.

## Supplemental Inventory

Supplementary materials

Table S1

Figure S1. **Hyaluronan production and degradation during implantation.** Related to Figures 1–2.

Figure S2. **Pharmacological inhibition of maternal hyaluronan production.**

## Supplementary material

**Table S1.** Viral titers measured for lentiviral vectors

| Insert | Vector | Viral titer (IFU/ml) |
| --- | --- | --- |
| Control | Lenti CRISPR v2 | $\geq 1 \times 10^7$ |
| HAS-1 del | Lenti CRISPR v2 HAS-1 deletion | $\geq 1 \times 10^7$ |
| HAS-2 del | Lenti CRISPR v2 HAS-2 deletion | $\geq 1 \times 10^7$ |

**Supplementary Figure 1.**
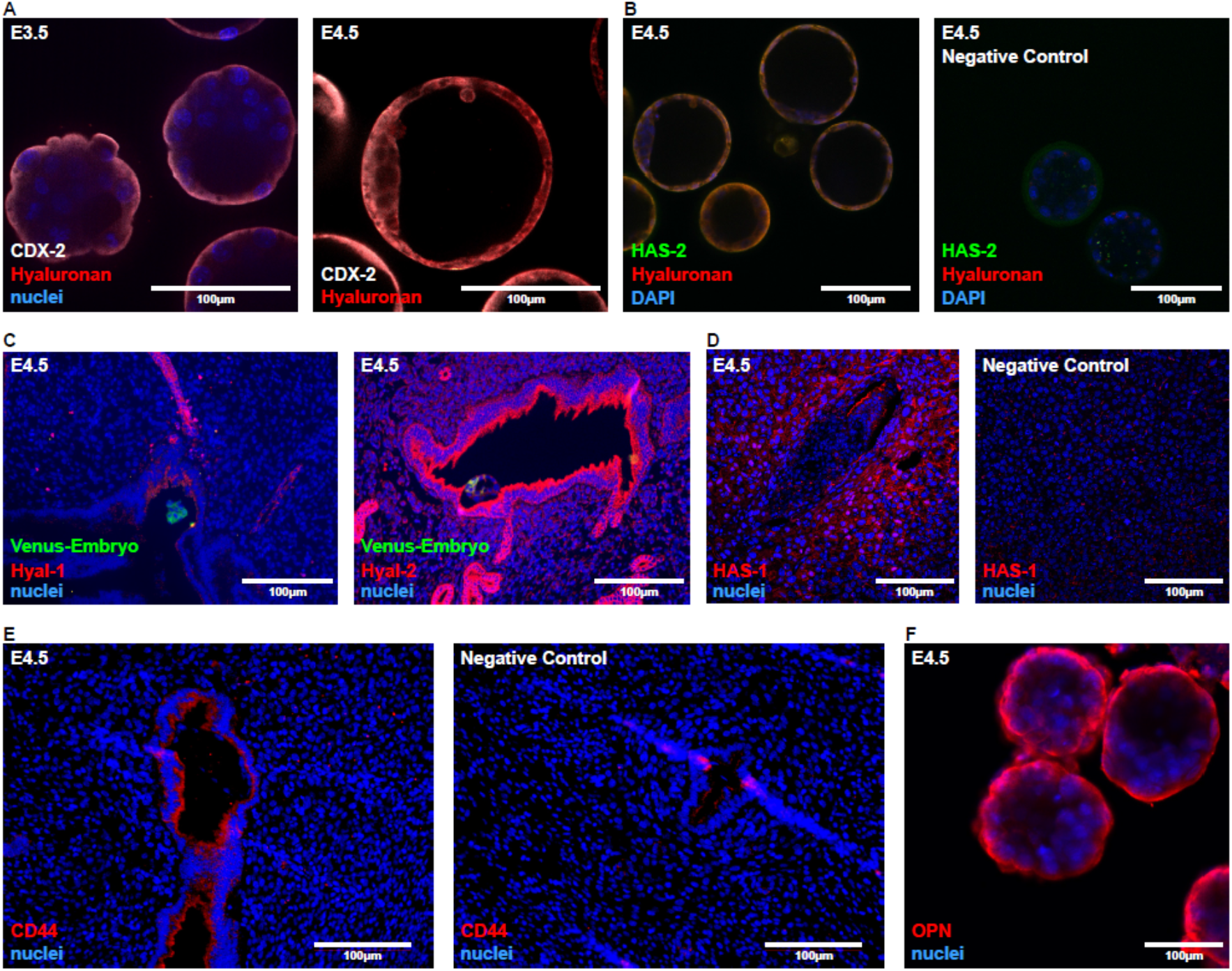
Hyaluronan production and degradation during implantation. Female mice were mated with Venus+ males, their uterine horns were harvested at E3.5 and E4.5 and wither immediately flushed or subjected to immunohostichenical analysis. (n=3 dams). (**A**) Immunofluorescent staining of Hyaluronan in CDX-2^+^ trophectoderm in blastocysts. (**B**) Immunofluorescent staining of hyaluronan and HAS-2 in blastocysts; Negative control was performed by excluding the primary antibody. (**C**) Immunohistochemical detection of Hyal-1 and Hyal-2 in histological sections. (**D**) Immunohistochemical detection of HAS-1 in histological sections at E6.5; Negative control was performed by excluding the primary antibody. (**E**) Immunohistochemical detection of CD44 in histological sections; Negative control was performed by excluding the primary antibody. (**F**) Immunofluorescent staining of osteopontin in pre-implantation blastocysts.

**Supplementary Figure 2.**
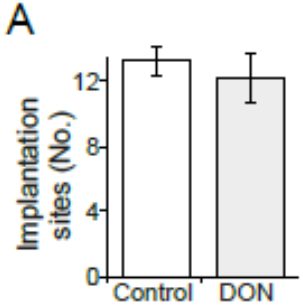
Pharmacological inhibition of maternal hyaluronan production. Pregnant ICR mice were administered with DON (1μg/g BW) daily at E3.5-E5.5 (**A**) Similar number of embryo implantation sites was observed both, for the control as well as for DON treated mice (13.14±0.82; 12.11±1.5) (n=9 dams in the control; 13 dams in DON-treated mice). (**B**) Surface area quantification of implantation sites (2.619 fold change±0.056; 0.037) (n=3 dams, 16 implantation sites in the control; 6 dams, 17 implantation sites in DON).

